# Social touch shapes communication and animal recognition in naked mole-rats

**DOI:** 10.1101/2024.02.21.581483

**Authors:** Ryan Schwark, Simon Ogundare, Preston Sheng, William Foster, Phalaen Chang, Yu-Young Tsai, Amanda Arnold, Ishmail Abdus-Saboor

## Abstract

The East African naked mole-rat (*Heterocephalus glaber*) lives in cooperative subterranean colonies and displays a capacity to recognize social novelty, rank, and identity. The sensory cues used for social recognition remain poorly understood, especially because many of their senses are either lost or greatly reduced in comparison to other mammals. Here, we found that naked mole-rats actively touch faces 100% of the time they encounter one another in a tunnel test, followed by milliseconds speed determination of the rank of the other animal. Even in an open arena, naked mole-rats engage in face-to-face touch hundreds of times in a 10-minute social pairing and colonies do so tens of thousands of times over a 24-hour period in their home environment. To demonstrate the prominence of face touch at a molecular level, we show that social housing conditions lead to widespread activation of mechanosensory ion channels, including Piezo2, in neurons that innervate the face, but not the body. Lastly, to determine the ethological relevance of face touch, we reduced its capacity with facial whisker trimming and revealed an apparent inability for animals to recognize colony members. Together, these findings uncover face touch as a prominent social behavior in naked mole-rats that is intimately linked to social recognition.

## Introduction

The naked mole-rat (*Heterocephalus glaber*) is one of the most social mammals in the animal kingdom (Buffenstein et al., 2021). These animals live in large colonies consisting of a breeding queen, 1-3 breeding males, and dozens to even hundreds of nonreproductive workers that are all descendants of the queen (Buffenstein et al., 2021; Jarvis, 1981). This close-knit and elaborate colony structure is stable over time, given that naked mole-rats live between 20-40 years (Buffenstein, 2005; Edrey et al., 2011; Ruby et al., 2018). Naked mole-rats engage in a range of collective behaviors including digging and expanding the colony, transporting food, and communal care of offspring (Buffenstein et al., 2021; Clarke & Faulkes, 1997; Watarai et al., 2018). The extreme sociability that naked mole-rats engage in is specific to colony members, as they do not allow foreign naked mole-rats entry into the colony. Thus, naked mole-rats have the ability to recognize their own colony members vs foreigners, and within their colonies, they are able to distinguish one animal from another (O’Riain & Jarvis, 1997). However, the mechanisms naked mole-rats use to recognize other naked mole-rats, and thus live long-term harmonious communal life, is largely unknown.

The sensory world of naked mole-rats is distinct from most above-ground animals, so the cues they use for social recognition might differ from other mammals. In terms of vocalization-audition, a recent study demonstrated that naked mole-rats generate dozens of unique vocalizations that are specific to a given colony, and the colony dialect is maintained by the presence of the queen (Barker et al., 2021; Pepper et al., 1991). In terms of smell, although olfaction is an important form of social communication for many rodents, the olfactory epithelia and bulb of naked mole-rats is greatly reduced compared to other mammals (Onyono et al., 2017), and their olfactory epithelium appears dispensable in a standard odor-based discrimination task (Toor et al., 2015). Although olfaction seems diminished in these animals, as well as vision (they are essentially blind due to millions of years of underground living) (Hetling et al., 2005), their sense of touch is quite prominent. In fact, although furless, naked mole-rats have facial vibrissae and vibrissa-like body hairs that are similar in morphology, and these vibrissae are exquisitely sensitive to deflections (Crish et al., 2003). A deflection of a single vibrissae on their body is sufficient to elicit a full body turn towards the axis of deflection – a somatosensory capacity that is unparalleled compared to most other mammals (Crish et al., 2003). Additionally, the somatosensory cortex of naked mole-rats is three-fold expanded compared to other mammals (Catania & Remple, 2002; Xiao et al., 2006), suggesting a heightened reliance on somatosensation. How somatosensation might be used to guide communication, enforce dominance, facilitate maintenance of social hierarchy, and support social recognition, is unknown.

Here, we study the social behaviors of naked mole-rats using automated video tracking coupled to behavior quantification at both regular and high-speed resolution. Intriguingly, we identify face-to-face contact as a prominent form of social interaction in naked mole-rats that appears indispensable for social recognition.

## Results

### Snout touch is a central component of dominance behavior in naked mole-rats

Mole-rat colonies can be divided into “ranks” based on dominance testing in which a more dominant mole-rat crawls over a more subordinate mole rat in a tube. To determine the dominance hierarchy within our colony, we used the tube test assay which is the standard in the field (Clarke & Faulkes, 1997, 1998; Hite et al., 2022; Toor et al., 2015). One observation that immediately caught our attention in this assay, that we will return to in greater detail below, is that face-to-face contact always preceded the rapid crawl-over decision (Figure 1A, left). A total of 18 mole-rats were used for every possible pairwise combination of animals, and with ten trials for each tube test, this amounted to ∼2,000 individual trials. From this data, overall crawl-over win% was used to construct a dominance hierarchy (Figure 1B) with higher win% animals assigned higher ranks. Worker win% ranged from less than 5% to over 85%, and the queen exhibited the second-highest win% and was assigned a rank of 2. It should be noted that the queen did not adopt the subordinate posture and “lose” tube tests; she would force herself underneath the other naked mole-rat (Figure S1). The relative ranks were then used for all subsequent experiments.

**Figure 1:**
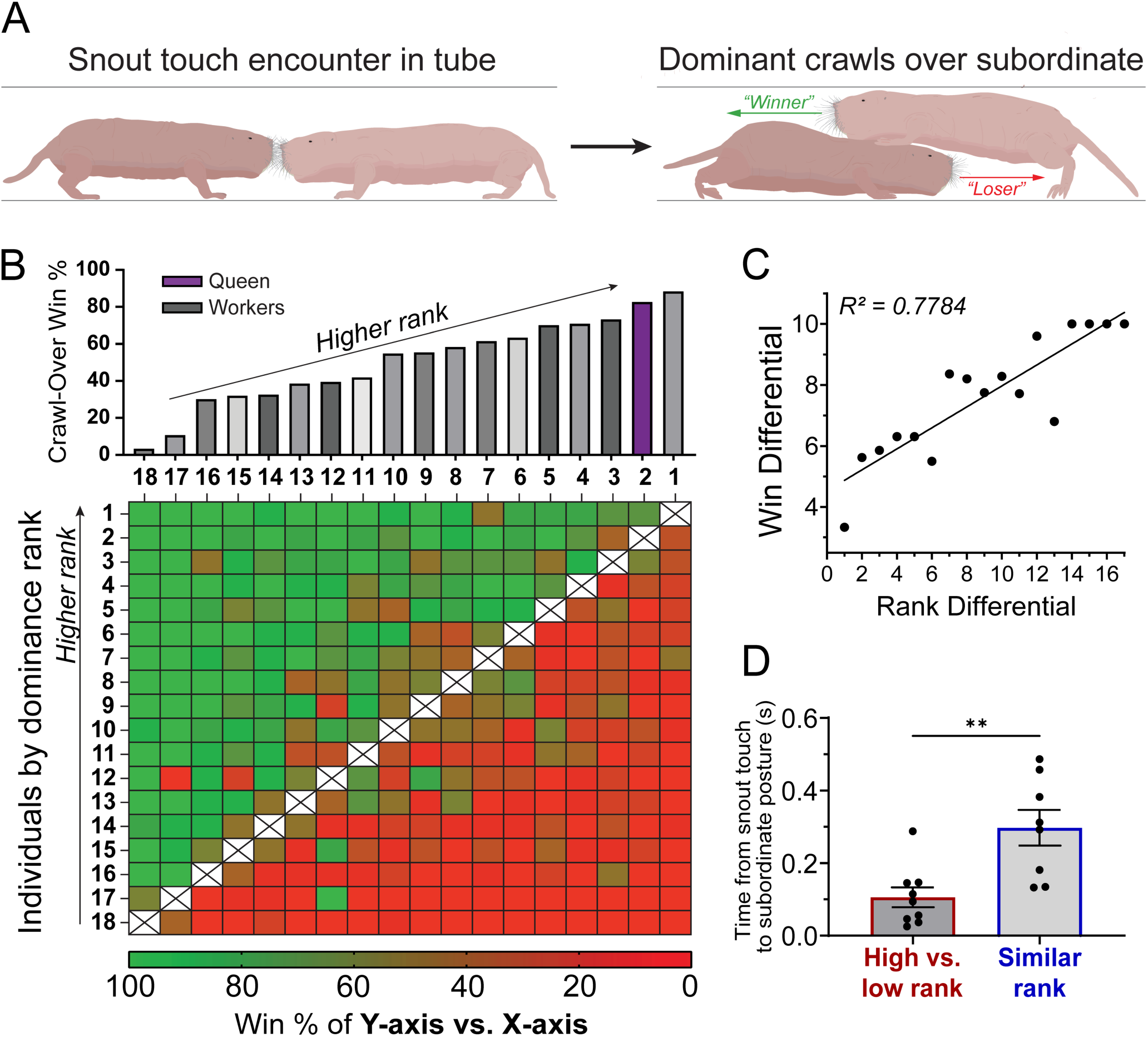
Snout touch is a central component of dominance behavior in naked mole-rats. **(A)** Schematic showing a single trial of the tube test for dominance in naked mole-rats. Upon encountering each other face-to-face in a tube, naked mole-rats will always engage in a snout interaction. Following this interaction, one naked mole-rat (the “winner”) crawls over the other naked mole-rat (the “loser”) which adopts a subordinate posture. **(B)** Dominance hierarchy of naked mole-rats generated through pairwise testing. Each pairwise tube test consisted of 10 consecutive trials, with color corresponding to win% of column Y vs. column X. Higher-ranked animals consistently “won” vs. lower-ranked individuals (n=18 animals). **(C)** Linear regression shows that the higher the rank differential between a pair of animals in the tube test, the greater the win differential is [R2 = 0.7784]. **(D)** Quantifying the time between snout touch and subordinate posture using high-speed videography. In pairings between animals with high differential in rank, latency is significantly faster than in animal pairings of similar rank [n = 9 tube tests for similar-rank pairs, 10 tests for high v. low rank pairs, two-tailed t-test, p=0.0350, Grubb’s test removed 2 outliers at alpha = 0.05].

As shown in Figure 1B, we then plotted the tube test results from every pairwise encounter in a matrix to determine how disparity in rank affects the outcome of each tube test. The results indicated that in 84.96% of encounters, the more dominant naked mole-rat won. Each encounter was assigned a rank differential, which corresponds to the distance between two naked mole-rats in the dominance hierarchy. Another metric which we created, win differential, quantified how lopsided the victory/loss of a given tube test was (win differential = |#wins_animal1_ – #wins_animal2_|). To determine how rank differential correlates with win differential, we plotted these two metrics (Figure 1C). The data showed that in a given tube test encounter, the greater the difference in rank between the two animals, the more asymmetrical the win is for the victorious animal (linear regression, R^2^ = 0.7784). This indicates that higher-ranked animals typically win, lower-ranked animals typically lose, and the win percentage predictably converges to 50% for evenly-ranked pairs.

Throughout these experiments, we noticed that snout-to-snout touch was quite common. We used high-speed videography to gain more clarity about these interactions (Figure 1D, Video S1). We found that snout touch interactions always occur first when two animals encounter each other, which is then followed by a crawl-over. When a high-dominance and a low-dominance animal encounter each other, the latency between snout touch and crawl-over is extremely fast (mean latency = 105.8 ms ± 0.0273 SEM). Comparatively, when the animals are similarly ranked, they spend a nearly tenfold greater time engaging in face touch (mean latency = 297.3 ms ± 0.0492 SEM) (Figure 1D). These results show that the speed of sensory processing during tube encounters occurs on very short timescales, and suggest that face touch might carry salient information about naked mole-rat identity. In other words, more closely ranked animals may need more time to gather information to determine the rank of their interlocutor in the tube.

### Active face touch is a prominent form of social interaction in naked mole-rats

We next quantified the extent of face touch interactions as animals freely interacted in a bucket using the pose estimation deep neural network SLEAP combined with our own custom code (see details in Methods) (Figure 2A; Video S3). We also compared naked mole-rat face touch interactions to mice as a reasonable rodent comparison. Remarkably, we discovered that naked mole-rats frequently socialize using active face touch, with an average of 100.8 interactions per 10 minutes (Figure 2B). This number of interactions was nearly double that of mice. This trend was also evident when quantifying the total duration of snout interactions over time (Figure 2C-D). The frequency of this behavior suggests its primacy in their social interactions.

**Figure 2:**
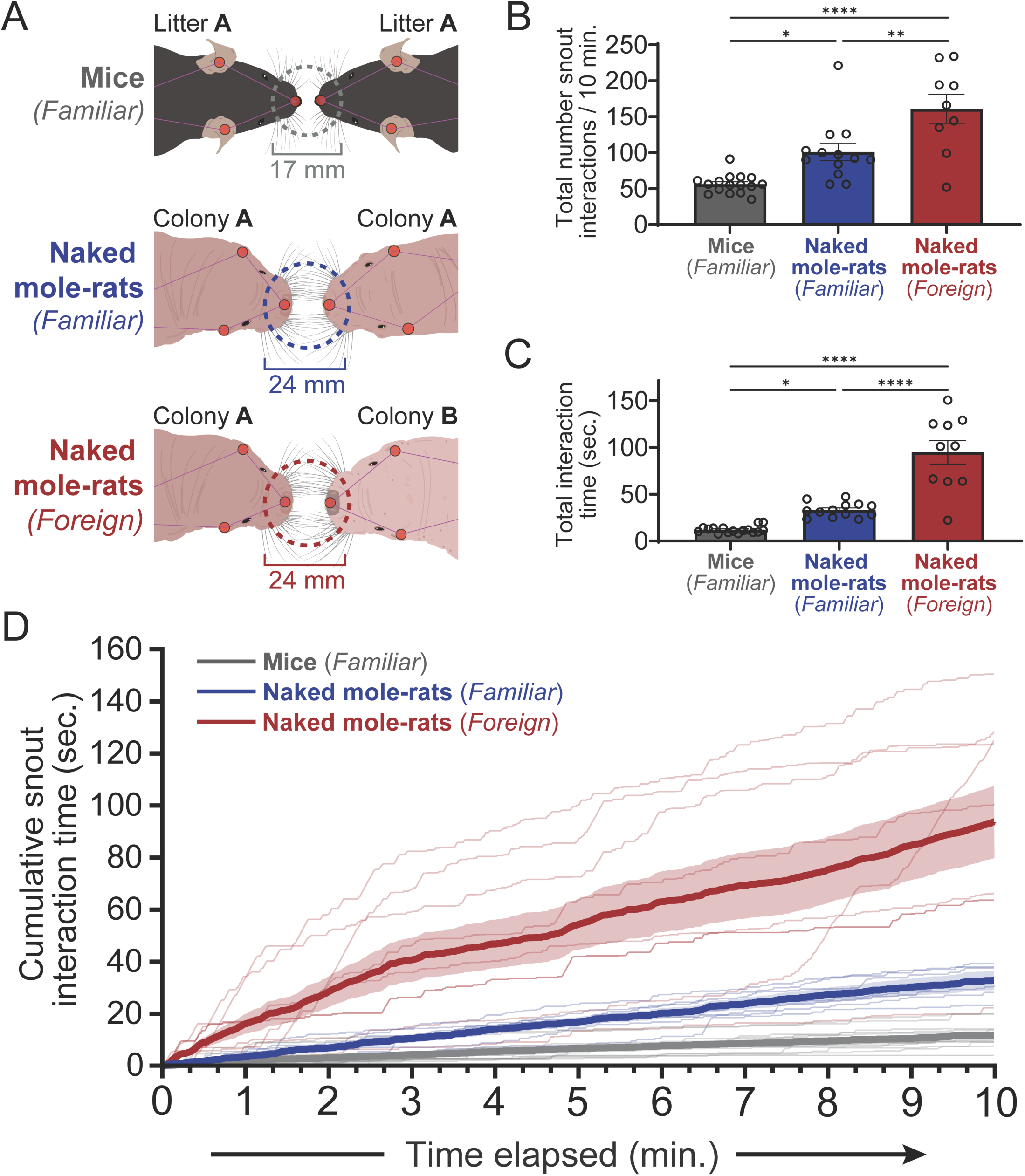
Face touch is a prominent form of social interaction in naked mole-rats. **(A)** Methods for tracking snout touch interactions in an open arena using SLEAP, comparing naked mole-rats to mice. A snout interaction between two naked mole-rats was defined as when snout tip proximity was 24 mm or less, determined by measuring the average distance at which the whiskers of the two animals could come into contact. The same methodology was used to determine a threshold of 17 mm in C57BL/6J mice. Open-field videos of the following pairs of animals were taken: co-housed familiar littermate mice (n=16), co-housed familiar littermate naked mole-rats from the same colony (n=12), and foreign naked mole-rats from separate colonies (n=9). **(B)** Number of snout interactions per 10 minutes. Two familiar naked mole-rats engage in more snout interactions (mean = 100.8 ± 3.26 interactions) than two familiar C57BL/6J mice (mean = 56.06 ± 11.67 interactions, One-Way ANOVA, p = 0.011). Two foreign naked mole-rats engage in substantially more snout interactions (mean = 161.0 ± 20.14 interactions) than familiar naked mole-rat pairs (One-Way ANOVA, p = 0.0030). **(C)** Total interaction time per 10 minutes of the same data in (B). Familiar naked mole-rats spend more time interacting (33.20 ± 0.934 seconds) than C57BL/6J mice (12.05 ± 2.16 seconds, One-Way ANOVA, p = 0.0234). Foreign naked mole-rats spend significantly more time interacting (94.71 05 ± 13.19 seconds) than familiar naked mole-rats (One-Way ANOVA, p < 0.0001). **(D)** Cumulative snout interaction time over a 10-minute period, comparing pairs of mice, familiar naked mole-rats, and foreign naked mole-rats.

Next, we decided to investigate face touch in the context of foreign encounters, since naked mole-rats are extremely xenophobic (O’Riain et al., 1996; O’Riain & Jarvis, 1997). For this experiment, we established additional naked mole-rat colonies donated from colleagues around the country. Interestingly, when we performed the experiments with foreign naked mole-rats in the same testing environment used with familiar animals (Figure 2B), we noticed a doubling of active face touch (Figure 2B,C). This result is consistent with our observations in a two-chamber preference assay where we show that naked mole-rats show increased interest in investigating a foreign naked mole-rat as opposed to a familiar conspecific (Figure S2). Thus, like mice, naked mole-rats appear to prefer novelty and have the memory capacity to remember members of their home colony, as well as members from other colonies. But why do foreign animals engage in even more active face touch? We speculate that the face touch is providing a cue or signal from one animal to another, and that relationship is altered amongst foreign animals, leaving the animals confused and spending more time trying to identify the other animal. We design an experiment later in the paper to test this hypothesis.

### Snout touch plays a prominent role inside the colony

To probe further into the ethological relevance of face touch interactions, we quantified snout touch in the colony setting where the naked mole-rats live. For these experiments, we developed a long-term monitoring system that continuously videotaped animals in their home colony (Figure 3A). Video was recorded from a digging chamber filled with a paper-based medium which the naked mole-rats dug through over time. Pose-tracking video from over a 24-hour period allowed us to quantify snout-to-snout interactions using the same 24 mm threshold as our previous experiments in the open field (Figure 3B). Strikingly, we found that snout-to-snout interactions were common inside the habitat, and exhibited a level of complexity not seen outside the home colony. Snout interaction magnitude—which corresponds to the number of naked mole-rat pairs engaging in a snout interaction at a given point in time—varied over the 24-hour recording session (Figure 3C). We found that the vast majority of snout interactions occurred between a pair of naked mole-rats, although more complex interactions also occurred (Figure 3D). Periods of higher activity were marked by snout interaction magnitudes of 2 and greater, reflecting multiple pairs of animals engaging in face touch at a given time. Furthermore, the majority of in-colony snout interactions were less than 2 seconds in length (Figure 3E). Taken together, we have identified active face touch in naked mole-rats as a prominent form of social interaction that occurs in both an open arena and during naturalistic social encounters in the home colony.

**Figure 3:**
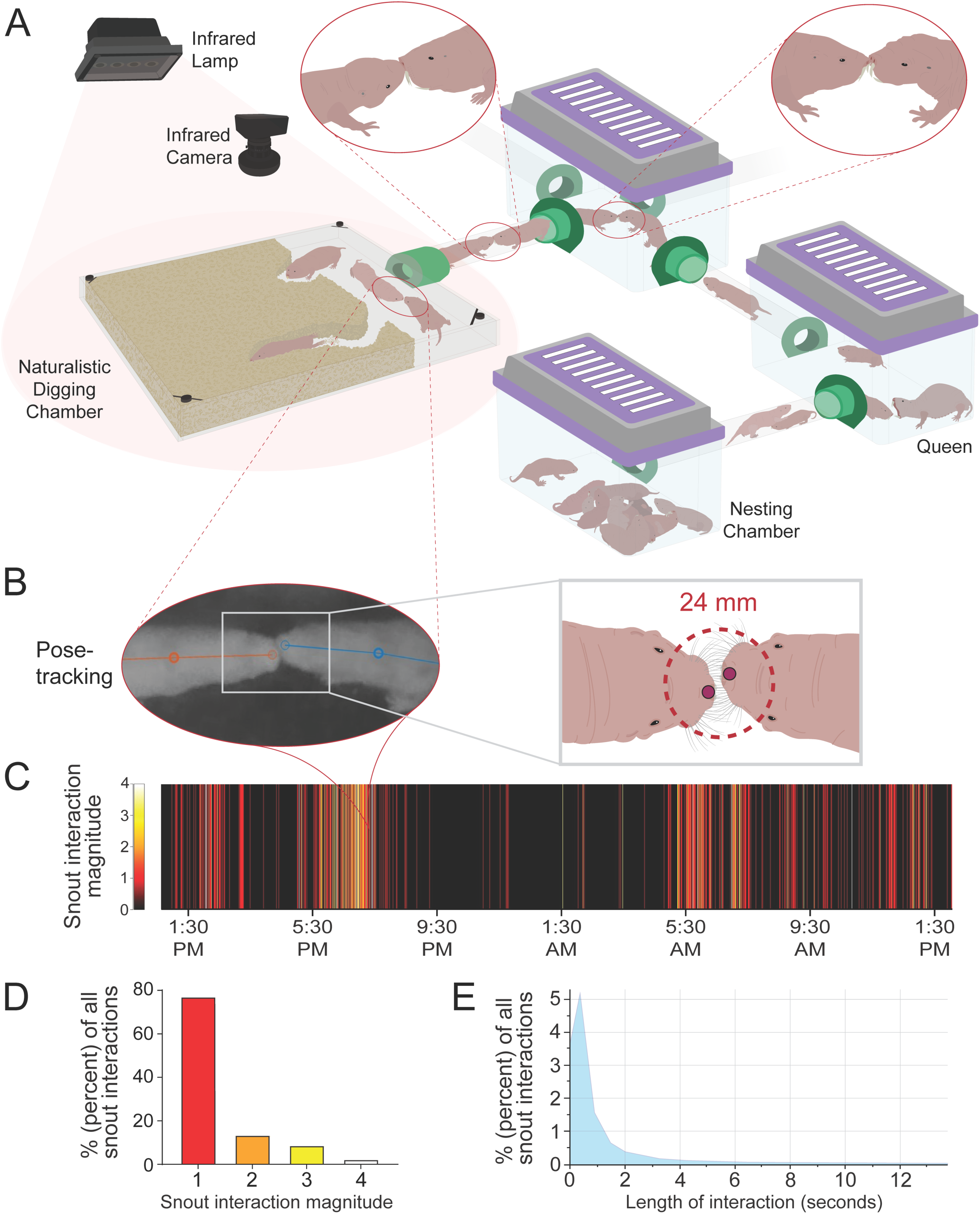
Snout touch is frequent and complex in the naked mole-rat home colony. (A) Schematic of long-term monitoring of snout touch in the naked mole-rat home colony. Animals frequently engage in snout touch throughout the interconnected colony, including in the naturalistic digging chamber. This digging chamber was continuously videotaped using an infrared camera, allowing snout interactions to be pose-tracked and quantified over a 24-hour period. (B) Pose-tracking and snout interaction quantification of naked mole-rats in the digging chamber. The threshold for a snout interaction was set at 24 mm, the same value as in the open-field experiments. (C) Extent of snout interactions in the digging chamber during a 24-hour period. Snout interaction magnitude corresponds to the number of two-animal pairs undergoing a snout interaction at a given time. Snout interaction magnitudes are color-coded, with darker regions corresponding to periods of low snout interaction in the digging chamber. (D) Distribution of snout interaction magnitudes during the 24-hour period in (C). The majority of all snout interactions were between two animals (75.31%). 13.15% of all snout interactions were of a magnitude 2, 8.50% were of magnitude 3, and 2.23% were of magnitude 4. (E) Distribution of snout interactions in the digging chamber by duration (seconds) The majority of snout interaction events lasted less than one second in duration; however, we also observed many interactions which were multiple seconds in length.

### Active face touch in social interactions involves mechanosensation in face-innervating sensory neurons with likely role of Piezo2

To determine the cellular basis of face touch within the home colony, we used the FM 1- 43 dye whose fluorescence intensity may be used as a readout of the opening and activation of mechanosensitive ion channels, including Piezo2, for example (Villarino et al., 2023) (Figure 4A). We tested either an isolated animal or an animal that remained in its home colony with ∼30 other naked mole-rats. The control and experimental animal were each given an intraperitoneal injection of FM 1-43 and were euthanized 24 hours later. In skin sections from snout skin of the face, within the social setting, we observed strong FM 1-43 fluorescence in lanceolate sensory terminal endings that associate with facial vibrissae (Figure 4B). These sensory endings in the face most likely emerge from Aβ mechanoreceptors, as these are the main mechanosensory endings in this snout region as previously described (Park et al., 2003). Strikingly, when we analyzed the trigeminal ganglion, we saw robust and widespread staining of the FM 1-43 dye in the social setting, but not when animals were alone, although in a similar home environment (Figure 4C, D). In the dorsal root ganglion, although we saw more FM 1-43 positive cells in the social setting vs. the isolated setting, the fluorescence intensity and number of fluorescent cells was greatly reduced compared to the trigeminal ganglion (Figure 4C-F). Since FM 1-43 can detect mechanosensitive ion channel activity beyond Piezo2, we wanted to confirm that naked mole-rats express Piezo2 in sensory neurons, evidence that would support the majority of FM 1-43 fluorescence intensity being linked to Piezo2 channel opening. Indeed, we showed that Piezo2 mRNA is present in both the TGs and DRGs using *in situ* hybridization (Figure 4H-I). Together, these results support the finding that active face touch, which is a highly mechanosensory behavior, is prominent when naked mole-rats socially engage. Moreover, naked mole-rats likely use the same molecular machinery for mechanosensation as other mammals despite being quite evolutionarily distant and divergent.

**Figure 4:**
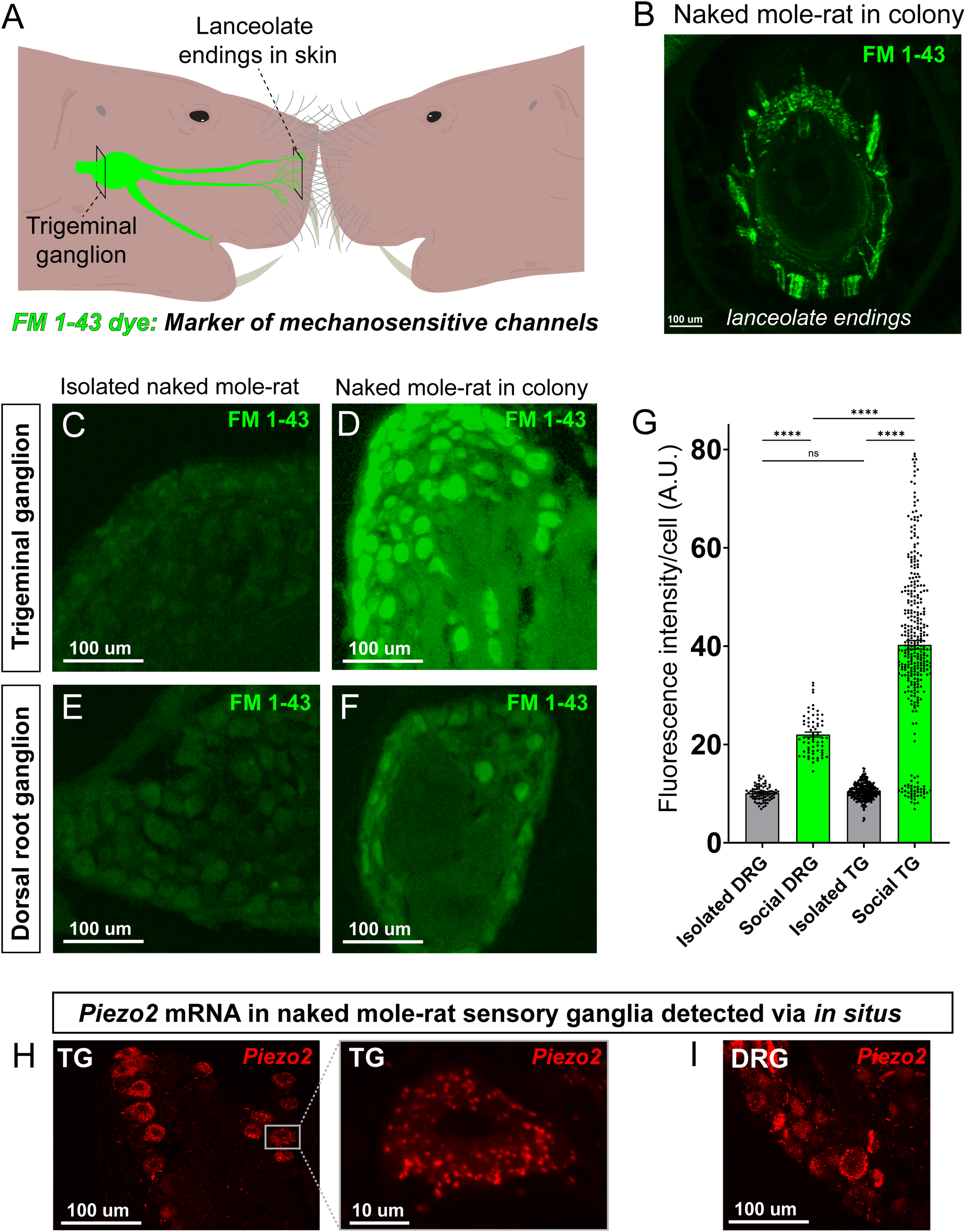
Face touch in the social setting involves activation of mechanosensitive channels such as Piezo2. **(A)** Schematic depicting peripheral innervation underlying face touch interaction in animals injected with mechanosensitive dye FM 1-43. Cryosections were obtained from the trigeminal ganglia innervating the face, and its peripheral sensory endings in the snout skin. **(B)** Activated lanceolate endings in the snout skin of naked mole-rat housed within social setting of home colony and injected with FM 1-43. Fluorescence indicates mechanosensitive activity in snout whiskers, with the majority of activity likely due to Piezo2. **(C)** Cryosection of trigeminal ganglion of naked mole-rat isolated from colony (and therefore isolated from snout-to-snout interactions) injected with FM 1-43. **(D)** Trigeminal ganglion section of FM 1-43-injected mole-rat housed in home colony and exposed to snout-to-snout interactions. **(E)** Dorsal root ganglion section of isolated naked mole-rat. **(F)** Dorsal root ganglion section of social naked mole-rat. **(G)** Comparison of peripheral neuron activation between naked mole-rats that were socially isolated vs. social (C-F). Quantifying fluorescence intensity per cell showed substantially higher Piezo2-dependent activity in the TGs and DRGs of social naked mole-rats than in socially isolated animals. In the social condition, the TG had substantially higher activity than the DRG [One-Way ANOVA, p<0.001, n=3 DRGs and 3 TGs per condition]. **(H)** *Piezo2* mRNA confirmed in naked mole-rat TGs using RNAScope protocol. Piezo2 transcripts could be seen in each individual cell (inset). **(I)** *Piezo2* mRNA was also detected in the DRGs, confirming expression of this mechanosensitive channel in the sensory neurons of naked mole-rats.

### Whisker trimming disrupts social dominance and animal recognition

In order to determine the role of the snout sensory whiskers in naked mole-rat face touch, we assayed how whisker trimming affected dominance behavior, and its connection to animal recognition, in the tube. Two cohorts of five naked mole-rats each were selected from the greater colony and their baseline dominance hierarchies and crawl-over latencies were determined. Strikingly, we found that when a whisker-trimmed naked mole-rat encountered a naked mole-rat with intact whiskers in the tube, crawl-over latency was significantly higher than at baseline (Figure 5A). Whereas the time from snout interaction to subordinate posture normally occurs on a timescale of hundreds of milliseconds, pairs with a whisker-trimmed animal took multiple seconds before initiating crawl-over. These results strongly suggest that the whiskers are crucial for individual recognition and that removing the whiskers causes animals to spend more time determining the identity of the other animal. Indeed, just watching the videos, it appears that the animals are repeatedly doing the face touch now, in a hopeless attempt to identify the other animal (Video S4).

**Figure 5:**
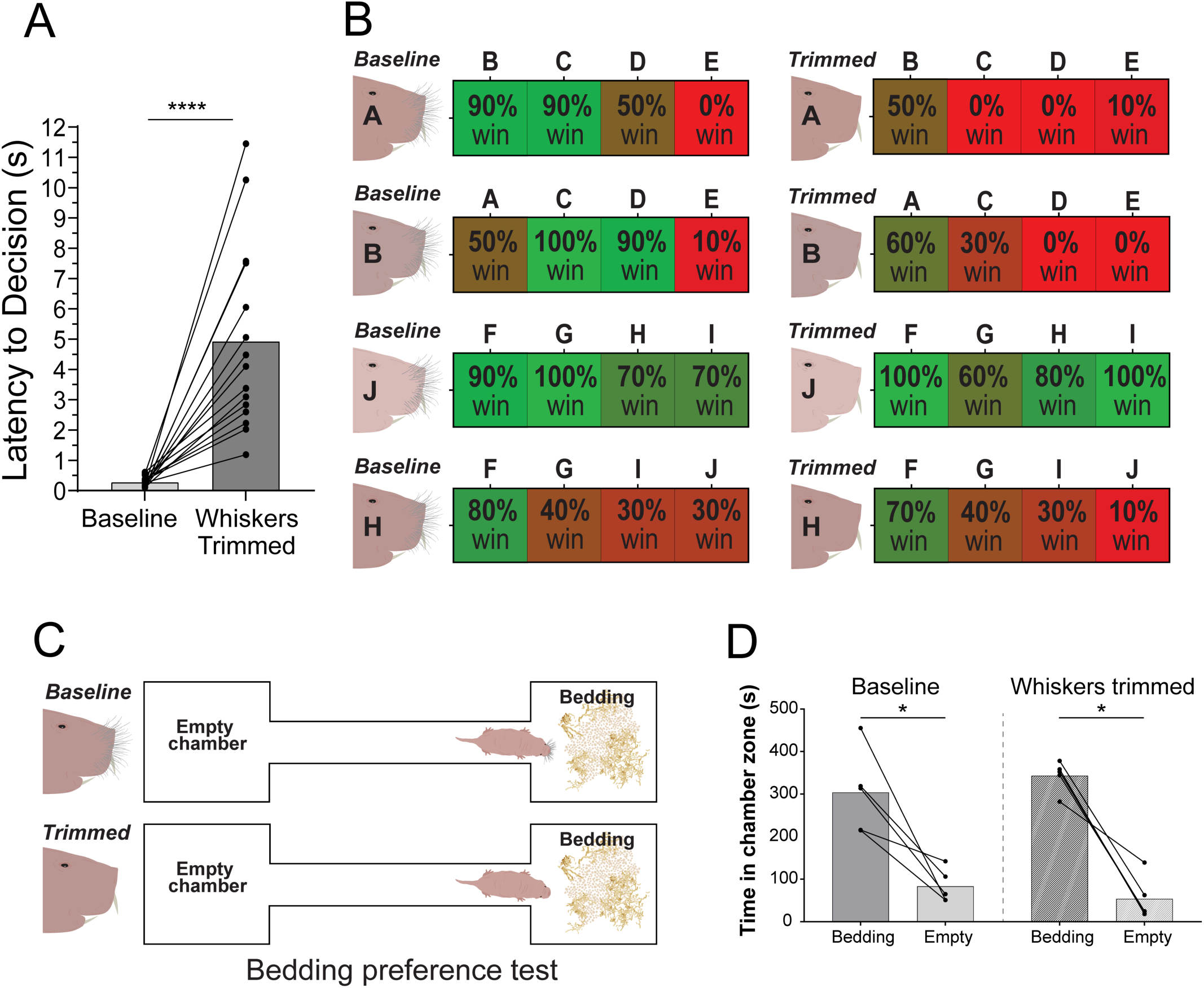
Trimming of the sensory whiskers increases tube test latency and disrupts dominance relationships. **(A)** Whisker trimming increases crawl-over latency during the tube test. Tube encounters between a whisker-trimmed animal and an animal with intact whiskers resulted in significantly longer snout interactions before crawl-over (mean = 4.920 ± 0.7928 seconds), compared to baseline (mean = 0.279 ± 0.0440 seconds). Latency was calculated as time between initial snout touch and adoption of subordinate posture by the “loser” in the tube test (One-way ANOVA, p<0.0001, n = 4 animals whisker trimmed). **(B)** Whisker trimming disrupts dominance relationships. Four animals were whisker trimmed (animals A, B, J and H). In all instances, the dominance relationships were altered. Whisker-trimmed animals often won fewer contests than at baseline, with many becoming more subordinate. Whisker trimming alternatively increased the win% for some pairings (paired two-tailed t-test, p = 0.0274, n=9 pairs for baseline and whisker trimmed conditions). **(C)** Schematic of bedding preference test. Baseline place preference for a chamber containing home colony bedding compared to an empty chamber was tested in naked mole-rats. Each animals’ preference was tested at baseline and after whisker trimming. **(D)** Olfactory-driven behavior remains intact after whisker trimming. Naked mole-rats exhibited a strong preference for home colony bedding at baseline with whiskers intact. Whisker trimmed animals retained this bedding preference, indicating that removing the whiskers does not broadly disrupt olfactory-driven behavior.

This deficit appeared to be present in both individuals—the naked mole-rat with intact whiskers may also have difficulty identifying the animal which lacks any whiskers. Relatedly, we discovered that removing the whiskers also disrupts the dominance relationships (Figure 5B). Whisker-trimmed individuals (animals A, B, J and H) exhibited significantly more subordinate behavior than at baseline (Figure 5B). As a control, we found that isoflurane anesthesia alone (which was used to trim the whiskers) was not sufficient to cause disruptions to crawl-over latency or dominance relationships (Figure S3). Moreover, the naked mole-rats still had a preference for home colony bedding in the absence of whiskers, demonstrating that overall behavior (especially non-somatosensory behavior) remains intact without whiskers (Figure 5C, D).

Taken together, these results suggests that removing the whiskers reduces naked mole-rats’ ability to recognize each other. Although the molecular and neural circuit basis of their whisker-based recognition system still remains to be discovered, these results demonstrate the ethological relevance of active face touch in naked mole-rat societies.

## Discussion

Although naked mole-rats have captivated biologists for decades, mainly for their longevity, resilience to cancer, and apparent indifference to certain types of pain (Buffenstein, 2008; Buffenstein et al., 2021; Freire Jorge et al., 2022), the biology of their social interactions has been less studied. Here, we establish face-to-face contact as a key social behavior—perhaps indispensable for the stability of the naked mole-rats’ reproductive and social hierarchies. Thus, these features of their biology and behavior make them an ideal model system for understanding the basis of complex social systems and social touch.

We observed naked mole-rats engaging in an active form of face touch hundreds of times over a short window. Moreover, in their home colony during naturalistic social behaviors, we observed tens of thousands of face touch interactions. Although we observed that mice also engage in snout-to-snout interactions, the frequency in naked mole-rats is much higher. Why do naked mole-rats engage in face touch so frequently? Perhaps the animals find this behavior rewarding, and checking in with other animals via touch provides social satiety, especially for a blind animal. However, our data is consistent with active face touch being part of their social communication and animal recognition system. Indeed, we observe an increased amount of face touch when two foreign naked mole-rats encounter one another—which might be due to increased efforts in trying to identify the stranger. The somatosensory system of the naked mole-rat is exquisitely tuned to respond to face touch by detecting movement on the skin through activation of vibrissae-innervating sensory neurons (Crish et al., 2006; Park et al., 2003). Our results here with the FM 1-43 dye in the trigeminal ganglion of naked mole-rats in a social setting is consistent with abundant face touch and direct mechanical activation of the somatosensory system. This result, coupled with *in situs* for *Piezo2*, shows that the naked mole-rat, separated by approximately 30 million years of evolution from the standard lab mouse, likely uses similar molecular machinery for mechanosensation. Removal of the whiskers from naked mole-rats significantly reduced their ability to recognize conspecifics and disrupted their dominance relationships, suggesting that the whiskers also play an important role in communication and identification. Moreover, the face, including the externally located incisor teeth, have a heightened representation in the somatosensory cortex compared to tactile inputs from other parts of the body (Catania & Remple, 2002). Studies in rats have shown that multiple cortical areas have their activity modulated by social snout-to-snout touch (Bobrov et al., 2014; Ebbesen et al., 2019) Thus, the naked mole-rat face is an ideal locus for gathering tactile sensory signals and transmitting this information to their expanded somatosensory cortex and other cortical and sub-cortical areas.

What about the role of other sensory systems in the social behavior of naked mole-rats? We cannot exclude the possibility that the close face contacts are made in an effort to smell a volatile compound or extract a pheromone cue on the other animal, or even to gain closeness to hear a unique vocalization. Indeed, active face touch seems unlikely as a sensory cue, in isolation, to be rich enough to serve as an animal identification code. A recent landmark study identified 25 unique vocalizations in naked mole-rat colonies (Barker et al., 2021), largely consistent with reports from an earlier study (Pepper et al., 1991). These vocalizations constitute a dialect that is culturally transmitted and maintained by the presence of a queen, and animals from the same colonies display vocal patterns of call and response (Barker et al., 2021). Indeed, spending any brief time amongst naked mole-rats, an investigator will easily discern an audible soft chirp. Multimodal communication using both vocalizations and whiskers has been shown in rats, suggesting that a similar mechanism may exist in naked mole-rats (Rao et al., 2014). To what degree naked mole-rats use these vocalizations as a system for individual identification or to guide social behaviors or social hierarchy remains unclear, especially since hearing capacity is diminished and auditory structures in the naked mole-rat outer and inner ear are greatly reduced (Heffner & Heffner, 1993; Mason et al., 2016; Okanoya et al., 2018). The role of olfaction in naked mole-rats remains controversial, with some studies showing olfactory-mediated colony preference and food localization (Judd & Sherman, 1996; O’Riain & Jarvis, 1997) and others studies showing no reduction in olfactory mediated behaviors when the olfactory epithelium is ablated (Toor et al., 2015). We hypothesize that animal recognition and social communication is multimodal, with varying contributions from olfaction, vocalization-audition, and somatosensation. How these sensory modalities integrate to drive social interactions in naked mole-rats is a fascinating question worthy of intense investigation.

## Methods

### Naked mole-rats husbandry and behavioral testing

All experimental testing was performed in compliance with the Guide for the Care and Use of Laboratory Animals (NIH). All procedures were approved by the Institutional Animal Care and Use Committee of Columbia University. Naked mole-rats were housed in a climate-controlled room maintained at 80-85° F and approximately 15-20% humidity, on a 12-hour day/light cycle. All experiments were conducted in this environment to ensure naturalistic conditions for the animals. Naked mole-rats were fed a daily diet of sweet potatoes, celery, and apple, and this diet was supplemented twice a week with ProNutro cereal enriched in vitamins (Bokomo, 100g Pronutro/16g protein mix). Naked mole-rat colonies were housed in interconnected habitats consisting of mouse and rat cages joined by plexiglass tubing, with the number of chambers determined by the size of the colony. For the colony gifted by Dr. Shelley Buffenstein, a specialized digging chamber was created and attached to the habitat. The digging chamber consisted of acrylic and had dimensions of 1” by 18” by 18”. The chamber was filled with a digging medium made of compacted newspaper. All naked mole-rats were microchipped using RFID transponders to track identity. Animals that were used for open field recording and dominance testing ranged from 3.09 to 12.34 years of age, and both sexes were equally tested. Breeding pairs were formed by housing a female and male naked mole-rat together in a rat cage. Naked mole-rats were generously provided by Dr. Rochelle Buffenstein, Dr. Dan McCloskey, and Dr. Thomas Park.

### Open field recording setup

For all open field recordings, naked mole-rats were recorded in a circular arena consisting of a linear low-density polyethylene (LLDPE) tank with a diameter of 17” (United States Plastic Corp., catalog #14317). The arena was filled with black terrarium sand (ZooMed ReptiSand, Midnight Black color) approximately ¼” deep to simulate a naturalistic digging medium for the animals. For even illumination, a 21-inch diameter ring light was positioned above the arena. Video on the arena was obtained using a BlackFly S USB3 FLIR camera (catalog #BFS-U3-13Y3M-C USB 3.1; 1.3 megapixels) with an 8mm UC Series Fixed Focal Length Lens (Edmund Optics; catalog #33-307). The camera was mounted on a tripod above the center of the arena, and videos were acquired at 60 FPS and 1280×1024 resolution. Videos were recorded using the Teledyne FLIR Spinnaker SDK SpinView software. For videos of isolated animals, naked mole-rats were placed in the center of the arena and recorded in segments of 10 minutes. 19 animals, including the queen, were videotaped for the SLEAP-to-MoSeq pipeline. For social contexts, naked mole-rats were introduced to the arena at opposite sides to allow for natural snout-to-snout encounters. The machines used for recording were running Windows 11 Pro, with a NVIDIA® GeForce RTX™ 3060 GDDR6 GPU, and an Intel® Core™ i7-11800H CPU processor.

### Pose estimation using SLEAP

Recorded videos were labeled using SLEAP, with a universal skeleton used for all naked mole-rats. This skeleton consisted of seven nodes: snout, left ear, right ear, centroid (torso), left hindpaw, right hindpaw, and base of the tail. All images were converted to grayscale upon import. We labeled 2,749 frames with which we then trained a model using the multi-animal top-down pipeline type. Our receptive field in the centroid model configuration was 76 pixels. Frames used to train this model were derived from videos of both isolated animals, videos with multiple animals, videos of male and female workers, and videos of the queen. The model was also trained on frames of animals with variable coloration to account for pigmentation variability in naked mole-rats. In order to run inference on every frame, we culled the max instances to the number of naked mole-rats in the respective video, enabled the “simple” cross-frame identity tracker method, and used the built-in functionality to connect single-track breaks. Any additional errors in tracking were corrected using the manual track proofreading function in SLEAP. All files were exported in HDF5 format for further analysis. The machine used to train the SLEAP model was running Windows 11 Pro, an NVIDIA GeForce RTX 3060 GPU, and an Intel Core i7-12700K CPU processor.

### Pairwise tube testing for dominance

For each pairwise tube test in the standard-sized tube, two naked mole-rats were removed from the colony and placed on opposite ends of a tube. A trial consisted of the two animals encountering each other in the tube, and concluded with one animal (the winner) crawling over the other (the loser). The next trial began by resetting the two naked mole-rats to their starting positions, and 10 trials were conducted for each pairwise combination of animals. 18 total naked mole-rats were tested from the colony provided by Dr. Rochelle Buffenstein with pairwise tests for every possible combination of animals (nCr = 153 combinations). For each pairwise test, the rank differential was calculated as the absolute value of the difference in rank (rank differential = |rank_animal1_ – rank_animal2_|). Another metric, win differential, was calculated as the absolute value of the difference in the number of wins (out of 10 trials) between the winner and loser naked mole-rat (win differential = |#wins_animal1_ – #wins_animal2_|). For some tube tests, a high-speed camera was used to capture the encounter at 750 - 2000 FPS with a Photron FastCAM Mini AX 50 170 K-M-32GB camera with a Zeiss 2/100M ZF.2 mount attachable lens. This camera was monochrome 170K, had 32 GB memory, and was controlled using a Dell laptop with Photron FastCAM software (PFV4). The time from first snout touch to adoption of subordinate posture was calculated using Photron PFV4. For experiments using naked mole-rats from the colony provided by Dr. Thomas Park, both the standard-diameter and small-diameter tubes were used. For small-diameter tube tests, animals could not crawl over each other due to size restrictions. As such, the winner of a given tube test was the animal which pushed the other animal out of the tube.

### Snout-to-snout interactions in open field

Snout interaction data was generated using custom code written in R (https://github.com/abdus-saboor-lab-code/NMR-Notebooks), applied to HDF5 files generated by SLEAP. This code extracted the x and y coordinates of each skeletal node. Using these coordinates, the distance between any two given nodes in pixels was calculated using the Pythagorean theorem =√((x2-x1)²+(y2-y1)²). The average whisker lengths of both naked mole-rats and C57BL/6J mice were measured using photographs next to a ruler. For each animal, the longest forward-facing whisker and the longest side-facing whisker were measured and average together to obtain an average maximum whisker length. The average maximum whisker length was calculated using 10 naked mole-rats, and 8 C57BL/6J mice. A snout interaction was defined as any continuous instance in which two snout nodes were within double the average maximum whisker length. In naked mole-rats, which were calculated to have an average maximum whisker length of 12 mm, this corresponded to a snout interaction threshold of ≤ 24 mm. For mice, which had an average maximum whisker length of 8.5 mm, the snout interaction threshold was ≤ 17 mm. Using these criteria, the code calculated the total number of snout interactions between all of the snout nodes in a given HDF5 file.

### Snout-to-snout interactions in the home colony

Videos of naked mole-rat behavior in the home colony were obtained from the previously-described digging chamber. An aluminum frame (McMaster-Carr, Cat#47065T101) built to surround an 18” by 18” by 1” digging chamber. Two infrared lights (ASIN B075F7NV56) were affixed to the top of the frame and illuminated the digging chamber. A White Matter e3 vision camera was positioned above the digging chamber and continuously acquired greyscale video at 15 FPS at 1080p. Videos were automatically saved as h264 mp4 files in 20-minute blocks. Videos were tracked using SLEAP, using a three-node skeleton consisting of the snout, centroid, and base of the tail. We labeled 1710 frames with which we then trained a model using the multi-animal top-down pipeline type. All files were exported in HDF5 format for further analysis. The machine used to train the SLEAP model was running Windows 11 Pro, an NVIDIA GeForce RTX 4090, and 13th Gen Intel(R) Core(TM) i9-13900 2.00 GHz processor. A snout interaction was defined as when distance between the snouts of two animals was less than a 24-millimeter threshold. We created a metric called “snout interaction magnitude” which corresponded to the total number of subthreshold snout pairs per video frame. The snout interaction magnitude was calculated for every frame of a 24-hour period within the digging chamber. From this dataset, the duration in time of each snout interaction magnitude was calculated.

### Social preference assay

Social preference assay was conducted in a custom chamber created from a polyethylene tank with dimensions of 12” width, 18” length, and 12” height (United States Plastic Corp., catalog# 14828). Two acrylic cage-like chambers were located at far corners of the bucket and held the naked mole-rats to serve as preference stimuli. The 10-minute trials were videotaped using the same BlackFly S USB3 FLIR camera as in the open field testing, and the investigating naked mole-rat was fully tracked using ANY-maze software (https://www.any-maze.com). The investigating naked mole-rat was placed in the bucket and was free to move and investigate either cage. During an initial habituation phase, the test subject was allowed to freely explore the paradigm for 15 minutes. Before each of the subsequent recording sessions, the social stimulus animal (i.e. in- and out-colony conspecifics) was placed in the respective chamber, which physically confined its movement and allowed limited interaction with the investigating animal through the barred fence. The test subject was then re-introduced to the center point of the field (opposite to the chamber location) at the start of each video recording. The chamber was disinfected in between trials.

### FM 1-43 injection and histology

In order to administer the mechanosensitive channel-detecting dye, naked mole-rats were scruffed and injected intraperitoneally (IP) with 1.12 mg/kg FM 1-43 (Thermo Fisher). For the socially isolated condition, the animal was removed from the home colony for 6 months and housed in a rat-sized cage within the same climate-controlled room as that of the colony. After 6 months, the animal was injected with FM 1-43. For the social condition, the injected naked mole-rat was housed in the home colony. Prior to tissue collection, both animals were perfused using 4% PFA, and tissue was harvested from trigeminal ganglia (TG), dorsal root ganglia (DRG), and snout skin. Tissues were fixed overnight in 4% PFA and then overnight in 30% sucrose in PBS. All tissues were then frozen in OCT until sectioning. For cryosectioning, TGs and DRGs were sectioned at 30 uM thickness, and snout skin was sectioned at 100 uM thickness, with cuts made parallel to the surface of skin. All tissues were mounted on SuperFrost microscope slides and imaged on a Leica confocal microscope. For TGs and DRGs, ImageJ was used to quantify fluorescence intensity and count cell bodies. The number of cells imaged from each condition were as follows; DRG (socially isolated), 83 cells; DRG (socially paired), 66 cells; TG (socially isolated), 273 cells; TG (socially paired), 334 cells.

### RNA in situ hybridization

Naked mole-rats were removed from the colony and perfused with PBS and 4% PFA. Tissue was harvested from the trigeminal ganglia (TG) and dorsal root ganglia (DRG) and fixed overnight in 4% PFA. This was followed by overnight immersion in 30% sucrose in PBS. TGs and DRGs were frozen in OCT and cryosectioned at 30uM thickness. Tissues were mounted on SuperFrost microscope slides and stored at −80° C. Slides were then subjected to the fixed frozen RNAScope *in situ* hybridization protocol (ACD). The probe used was for Piezo2 mRNA (ACD, Catalog #400191) and the fluorophore used to visualize transcripts was Opal 570. Slices were imaged using a confocal microscope (Nikon A1R). ImageJ was used to extract representative images from z-stacks. Tissues were obtained from 2 naked mole-rats.

### Whisker trimming experiments

Prior to whisker trimming, two cohorts of five naked mole-rats each were selected from a larger colony. The baseline dominance hierarchy of each cohort was established via tube testing, with 10 trials per pair using a standard-sized tube. Baseline crawl-over latencies were quantified as previously described. Before an animal was whisker trimmed, it was subjected to an isoflurane control condition in which the naked mole-rat was anesthetized and allowed to recover is isolation for 3 hours. After isoflurane anesthesia, the naked mole-rat was run through identical tube testing procedures as before to quantify the effects of anesthesia on dominance behavior. The following day, the naked mole-rat was anesthetized using isoflurane under identical conditions as before and the whiskers were trimmed to the base using surgical scissors. After a 3-hour recovery period, the whisker trimmed naked mole-rat was run through dominance testing within its respective cohort of 4 other naked mole-rats.

### Bedding preference assay

For the bedding preference assay, a custom two-chamber device was constructed out of acrylic. Each chamber (left and right) was of dimensions 6” width by 6.5” length, and the two chambers were connected by a narrow passage of 12” length and 2” width. For baseline testing, a naked mole-rat with intact whiskers was placed in the center of the narrow passage, at the midpoint between both chambers. One chamber was left empty, while the other chamber was filled with familiar bedding from the naked mole-rats’ home colony. Each naked mole-rat was allowed to explore the chambers for 10 minutes. Each of the bedding preference trials were videotaped using a BlackFly S USB3 FLIR camera as in the open field testing. Animals were fully tracked using SLEAP, and the amount of time the animal spent in each of the two chambers was calculated. Five total naked mole-rats were tested. These same five animals were then whisker trimmed as previously described, and subsequently run through the same bedding preference assay as before.

## Data availability

All data and code are available upon request.

## Acknowledgements

We thank members of the Abdus-Saboor lab as well as Richard Axel, Larry Abbott, and Scott Linderman for helpful discussion and comments on this study and manuscript. We are indebted to Thomas Park, Ewan St. John Smith, Dan McCloskey, and Shelley Buffenstein for generously sharing naked mole-rats, as well as information on husbandry and best practices. We thank members of Ryan’s thesis committee, Stuart Firestein and Erin Barnhart, for helpful suggestions. IA-S and lab members acknowledge support from Columbia University start-up funds, Howard Hughes Medical Institute, National Institute of Health New Innovator Award, Rita Allen Foundation, Pew Charitable Trust, Brain Research Foundation, McKnight Foundation, Burroughs Wellcome Fund, Simons Foundation, Alfred P. Sloan Foundation, and Chan Zuckerberg Initiative.

---

**Supplemental Figure 1:**
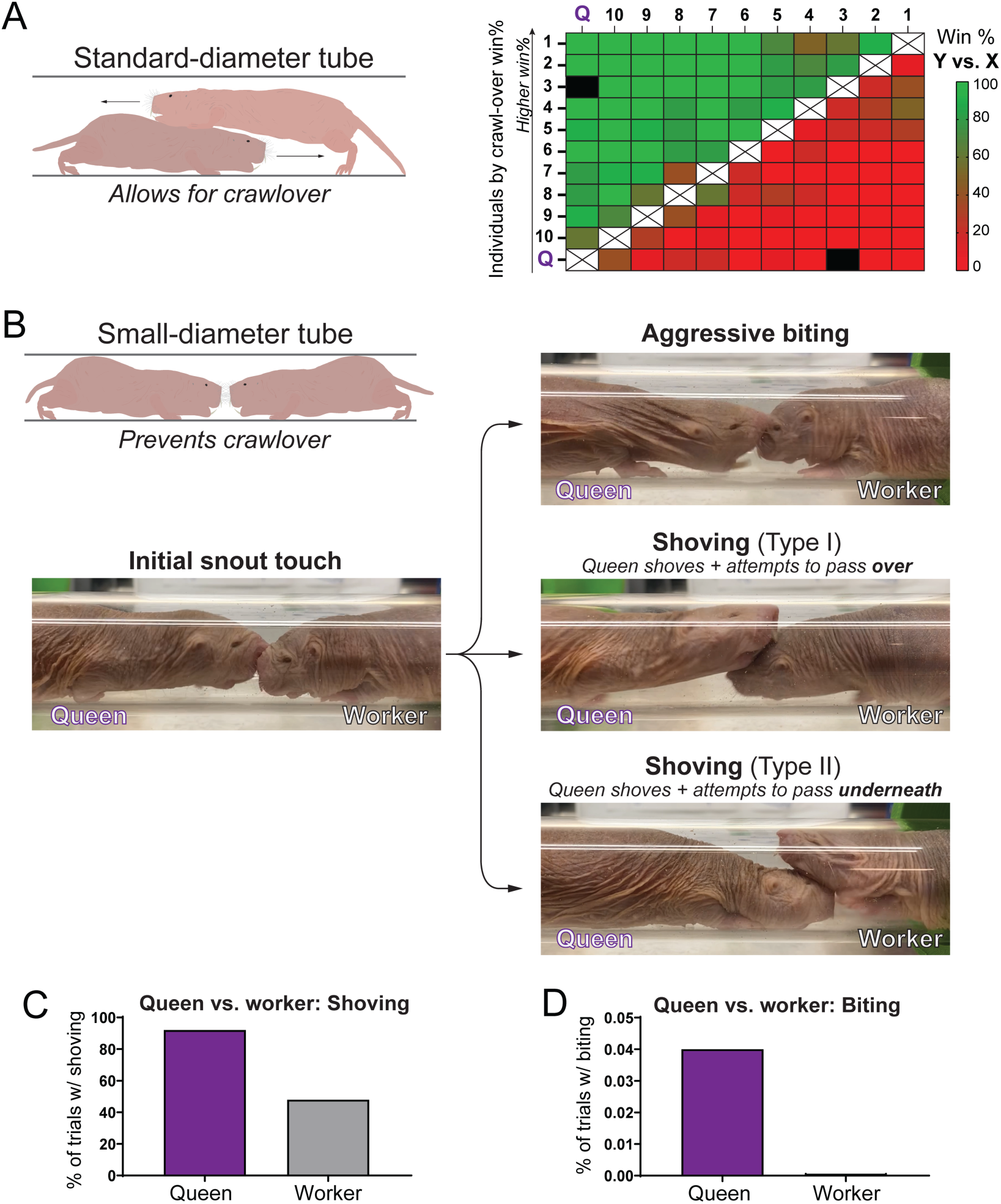
Tube testing results are consistent across colonies but the queen does not follow the rules of “etiquette” in the tube. **(A)** Determining the dominance hierarchy of a colony obtained from the Park lab. Using a standard-diameter tube which allows for crawl-over, the pairwise heat map exhibited an overall similar pattern to that of the Abdus-Saboor colony. One notable exception was that the Park colony queen crawled under during the vast majority of tests, leading to her ranked last by win%. **(B-D)** Testing the Park colony queen with a small-diameter tube reveals that she exhibits dominant crawl-under behavior and does not lose tube tests in the traditional sense. The queen frequently showed aggressive behavior and shoving of workers, and did not exhibit the arched back characteristic of subordinate capitulation. The queen frequently shoved underneath the workers, resulting in her “losing” often. This aggressive queen-specific phenotype suggests naked mole-rat queens may abide by a different set of tube test rules than the workers. n= 10 queen-worker trials, 3 different workers tested.

**Supplemental Figure 2:**
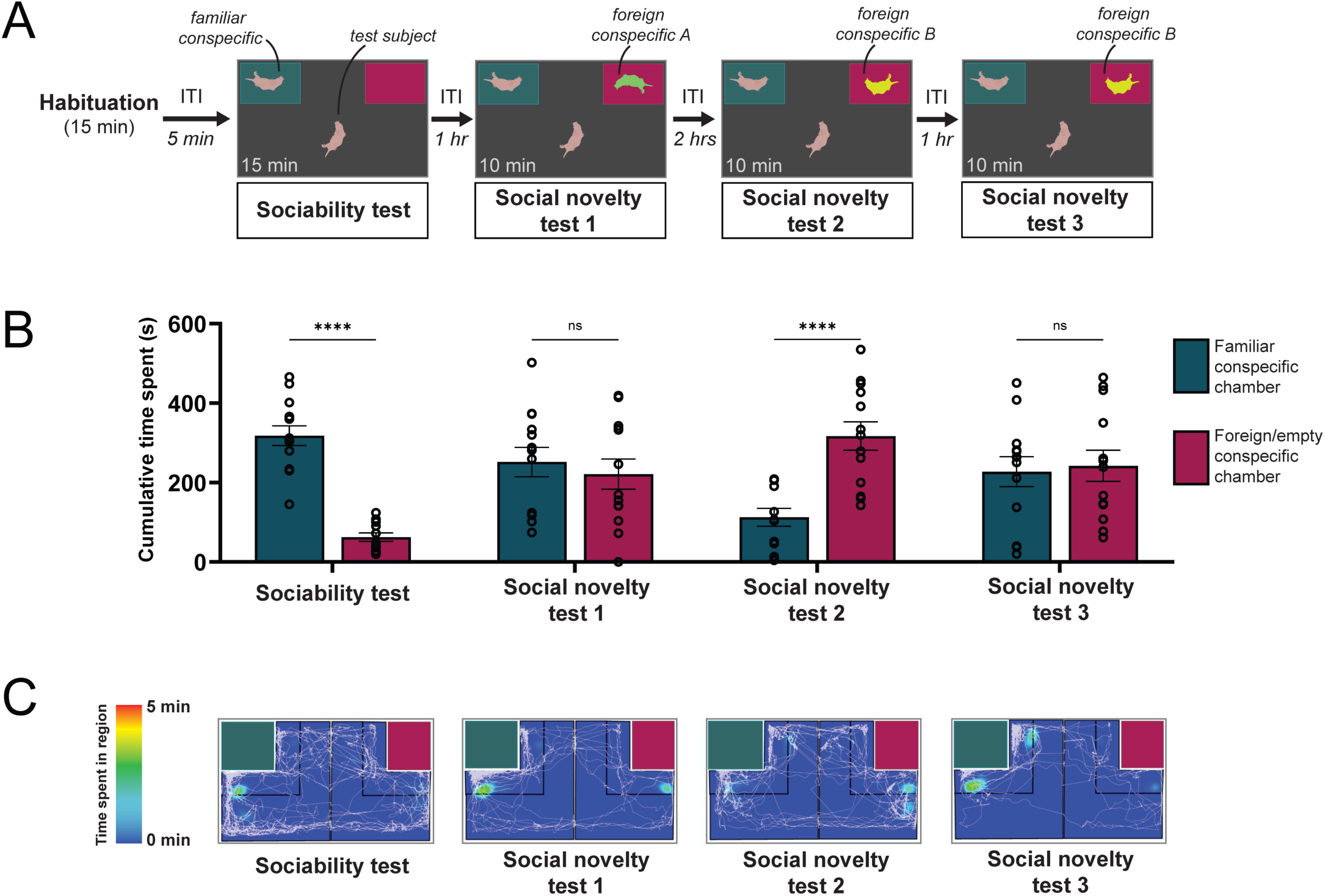
Naked mole-rats exhibit strong place preference for conspecifics. **(A)** Experimental design of social preference assay. After a 15-minute habituation period to two empty chambers and a 5-minute inter-trial interval (ITI), the test subject naked mole-rat underwent a sociability test with a familiar conspecific vs. an empty chamber for 15 minutes. Three subsequent social novelty tests then allowed the test animal to choose between a familiar vs. a foreign conspecific from another colony. **(B)** Naked mole-rats exhibit strong social preference and show signs of social memory towards foreign animals. Investigating animals overwhelmingly prefer to investigate a familiar conspecific vs. an empty chamber and spend similar time investigating a familiar vs. foreign conspecific. During social novelty test 2, preference for the foreign conspecific is higher, but this preference is lost during the subsequent social novelty test 3. This loss of preference suggests habitation to the foreign conspecific, potentially due to social memory. **(C)** Representative heat map of investigating animal during each test condition. Color indicates areas of density of the animal’s centroid over time. White lines indicate the trajectory of the snout over time [n=13 test subject naked mole-rats and respective total trials].

**Supplemental Figure 3:**
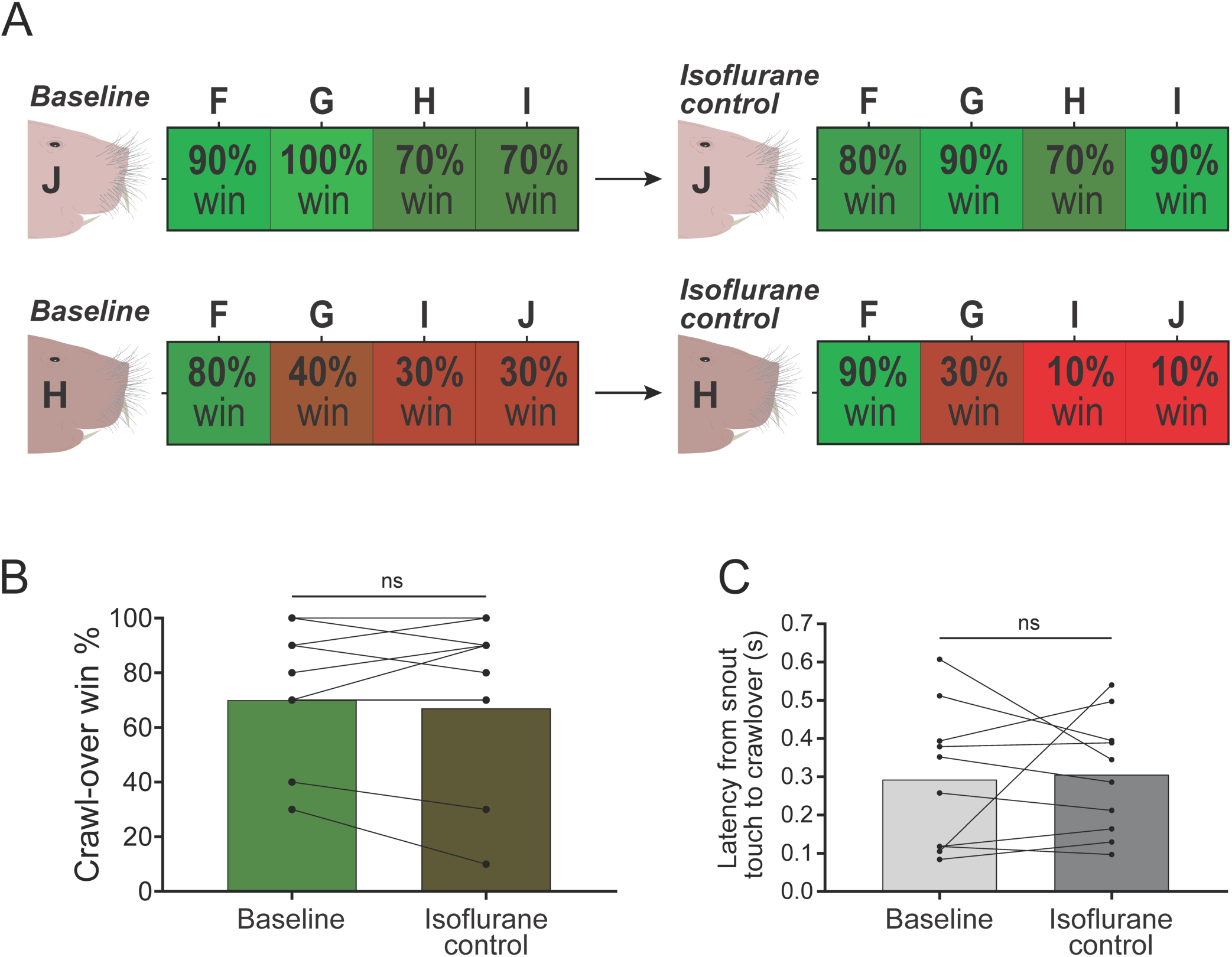
Isoflurane anesthesia does not affect social behavior in the tube test. **(A)** Isoflurane anesthesia does not affect dominance relationships. Anesthetizing naked mole-rats with isoflurane, and allowing them to recover, does not result in noticeable changes to dominance relationships in the tube test. **(B)** Quantification of the data in (A), comparing crawl-over win% between isoflurane-anesthetized naked mole-rats and baseline. No significant changes to crawl-over win% was seen between the two groups (paired two-tailed t-test, α = 0.05, p=0.4961, n=10 pairs). **(C)** Isoflurane anesthesia has no effect on crawl-over latency in the tube test (paired two-tailed t-test, α = 0.05, p=0.8258, n=10 pairs).

**Supplemental Video 1: High-speed video of tube test for dominance.** Sample of two naked mole-rats of similar rank during the tube test, as described in Figure 2E. The interaction begins with a snout-to-snout touch, followed by adoption of subordinate posture by the lower-rank animal, and crawl-over of the higher-rank animal. Video was recorded at framerate of 750FPS.

**Supplemental Video 2: Open field SLEAP tracking of two naked mole-rats.** Sample of two naked mole-rats fully identity-tracked by SLEAP, using the methodology described in Figure 1A. One animal (blue skeleton) engages in digging behavior, and the other (orange skeleton) explores the arena.

**Supplemental Video 3: Snout interaction of two familiar naked mole-rats.** Sample of snout interactions between two naked mole-rats from the same colony, as described in Figure 3A. Two snout interactions occur in this video, and prominently involve face touch.

**Supplemental Video 4: Whisker trimming increases tube test crawl-over latency.** Sample video comparing crawl-over latency between two naked mole-rats at baseline with intact whiskers, to the same pairing after one of the animals was whisker trimmed.

## References

Barker, A. J., Veviurko, G., Bennett, N. C., Hart, D. W., Mograby, L., & Lewin, G. R. (2021). Cultural transmission of vocal dialect in the naked mole-rat. Science, 371(6528), 503–507. 10.1126/science.abc6588

Bobrov, E., Wolfe, J., Rao, R. P., & Brecht, M. (2014). The Representation of Social Facial Touch in Rat Barrel Cortex. Current Biology, 24(1), 109–115. 10.1016/j.cub.2013.11.049

Buffenstein, R. (2005). The Naked Mole-Rat: A New Long-Living Model for Human Aging Research. The Journals of Gerontology Series A: Biological Sciences and Medical Sciences, 60(11), 1369–1377. 10.1093/gerona/60.11.1369

Buffenstein, R. (2008). Negligible senescence in the longest living rodent, the naked mole-rat: Insights from a successfully aging species. Journal of Comparative Physiology B, 178(4), 439–445. 10.1007/s00360-007-0237-5

Buffenstein, R., Park, T. J., & Holmes, M. M. (Eds.). (2021). The Extraordinary Biology of the Naked Mole-Rat (Vol. 1319). Springer International Publishing. 10.1007/978-3-030-65943-1

Catania, K. C., & Remple, M. S. (2002). Somatosensory cortex dominated by the representation of teeth in the naked mole-rat brain. Proceedings of the National Academy of Sciences, 99(8), 5692–5697. 10.1073/pnas.072097999

Clarke, F. M., & Faulkes, C. G. (1997). Dominance and queen succession in captive colonies of the eusocial naked mole–rat, Heterocephalus glaber. Proceedings of the Royal Society of London. Series B: Biological Sciences, 264(1384), 993–1000. 10.1098/rspb.1997.0137

Clarke, F. M., & Faulkes, C. G. (1998). Hormonal and behavioural correlates of male dominance and reproductive status in captive colonies of the naked mole–rat, Heterocephalus glaber. Proceedings of the Royal Society of London. Series B: Biological Sciences, 265(1404), 1391–1399. 10.1098/rspb.1998.0447

Crish, S. D., Dengler-Crish, C. M., & Comer, C. M. (2006). Population coding strategies and involvement of the superior colliculus in the tactile orienting behavior of naked mole-rats. Neuroscience, 139(4), 1461–1466. 10.1016/j.neuroscience.2005.11.073

Crish, S. D., Rice, F. L., Park, T. J., & Comer, C. M. (2003). Somatosensory Organization and Behavior in Naked Mole-Rats I: Vibrissa-Like Body Hairs Comprise a Sensory Array That Mediates Orientation to Tactile Stimuli. Brain, Behavior and Evolution, 62(3), 141–151. 10.1159/000072723

Ebbesen, C. L., Bobrov, E., Rao, R. P., & Brecht, M. (2019). Highly structured, partner-sex- and subject-sex-dependent cortical responses during social facial touch. Nature Communications, 10(1), 4634. 10.1038/s41467-019-12511-z

Edrey, Y. H., Hanes, M., Pinto, M., Mele, J., & Buffenstein, R. (2011). Successful Aging and Sustained Good Health in the Naked Mole Rat: A Long-Lived Mammalian Model for Biogerontology and Biomedical Research. ILAR Journal, 52(1), 41–53. 10.1093/ilar.52.1.41

Freire Jorge, P., Goodwin, M. L., Renes, M. H., Nijsten, M. W., & Pamenter, M. (2022). Low Cancer Incidence in Naked Mole-Rats May Be Related to Their Inability to Express the Warburg Effect. Frontiers in Physiology, 13, 859820. 10.3389/fphys.2022.859820

Heffner, R. S., & Heffner, H. E. (1993). Degenerate hearing and sound localization in naked mole rats (Heterocephalus glaber), with an overview of central auditory structures. Journal of Comparative Neurology, 331(3), 418–433. 10.1002/cne.903310311

Hetling, J. R., Baig-Silva, M. S., Comer, C. M., Pardue, M. T., Samaan, D. Y., Qtaishat, N. M., Pepperberg, D. R., & Park, T. J. (2005). Features of visual function in the naked mole-rat Heterocephalus glaber. Journal of Comparative Physiology A, 191(4), 317–330. 10.1007/s00359-004-0584-6

Hite, N. J., Sudheimer, K. D., Anderson, L., & Sarko, D. K. (2022). Spatial Learning and Memory in the Naked Mole-Rat: Evolutionary Adaptations to a Subterranean Niche. Frontiers in Ecology and Evolution, 10, 879989. 10.3389/fevo.2022.879989

Jarvis, J. U. M. (1981). Eusociality in a Mammal: Cooperative Breeding in Naked Mole-Rat Colonies. Science, 212(4494), 571–573. 10.1126/science.7209555

Judd, T. M., & Sherman, P. W. (1996). Naked mole-rats recruit colony mates to food sources. Animal Behaviour, 52(5), 957–969. 10.1006/anbe.1996.0244

Mason, M. J., Cornwall, H. L., & Smith, E. St. J. (2016). Ear Structures of the Naked Mole-Rat, Heterocephalus glaber, and Its Relatives (Rodentia: Bathyergidae). PLOS ONE, 11(12), e0167079. 10.1371/journal.pone.0167079

Okanoya, K., Yosida, S., Barone, C. M., Applegate, D. T., Brittan-Powell, E. F., Dooling, R. J., & Park, T. J. (2018). Auditory-vocal coupling in the naked mole-rat, a mammal with poor auditory thresholds. Journal of Comparative Physiology A, 204(11), 905–914. 10.1007/s00359-018-1287-8

Onyono, P. N., Kavoi, B. M., Kiama, S. G., & Makanya, A. N. (2017). Functional Morphology of the Olfactory Mucosa and Olfactory Bulb in Fossorial Rodents: The East African Root Rat (Tachyoryctes splendens) and the Naked Mole Rat (Heterocephalus glaber). Tissue and Cell, 49(5), 612–621. 10.1016/j.tice.2017.07.005

O’Riain, M. J., & Jarvis, J. U. M. (1997). Colony member recognition and xenophobia in the naked mole-rat. Animal Behaviour, 53(3), 487–498. 10.1006/anbe.1996.0299

O’Riain, M. J., Jarvis, J. U. M., & Faulkes, C. G. (1996). A dispersive morph in the naked mole-rat. Nature, 380(6575), 619–621. 10.1038/380619a0

Park, T. J., Comer, C., Carol, A., Lu, Y., Hong, H. -S., & Rice, F. L. (2003). Somatosensory organization and behavior in naked mole-rats: II. Peripheral structures, innervation, and selective lack of neuropeptides associated with thermoregulation and pain. Journal of Comparative Neurology, 465(1), 104–120. 10.1002/cne.10824

Pepper, J., Braude, S. H., Lacey, E., & Sherman, P. (1991). 9. Vocalizations of the Naked Mole-Rat. In P. W. Sherman, J. U. M. Jarvis, & R. D. Alexander (Eds.), The Biology of the Naked Mole-Rat (pp. 209– 242). Princeton University Press. 10.1515/9781400887132-011

Rao, R. P., Mielke, F., Bobrov, E., & Brecht, M. (2014). Vocalization–whisking coordination and multisensory integration of social signals in rat auditory cortex. eLife, 3, e03185. 10.7554/eLife.03185

Ruby, J. G., Smith, M., & Buffenstein, R. (2018). Naked mole-rat mortality rates defy Gompertzian laws by not increasing with age. eLife, 7, e31157. 10.7554/eLife.31157

Toor, I., Clement, D., Carlson, E. N., & Holmes, M. M. (2015). Olfaction and social cognition in eusocial naked mole-rats, Heterocephalus glaber. Animal Behaviour, 107, 175–181. 10.1016/j.anbehav.2015.06.015

Villarino, N. W., Hamed, Y. M. F., Ghosh, B., Dubin, A. E., Lewis, A. H., Odem, M. A., Loud, M. C., Wang, Y., Servin-Vences, M. R., Patapoutian, A., & Marshall, K. L. (2023). Labeling PIEZO2 activity in the peripheral nervous system. Neuron, 111(16), 2488–2501.e8. 10.1016/j.neuron.2023.05.015

Watarai, A., Arai, N., Miyawaki, S., Okano, H., Miura, K., Mogi, K., & Kikusui, T. (2018). Responses to pup vocalizations in subordinate naked mole-rats are induced by estradiol ingested through coprophagy of queen’s feces. Proceedings of the National Academy of Sciences, 115(37), 9264– 9269. 10.1073/pnas.1720530115

Xiao, J., Levitt, J. B., & Buffenstein, R. (2006). A stereotaxic atlas of the brain of the naked mole-rat (Heterocephalus glaber). Neuroscience, 141(3), 1415–1435. 10.1016/j.neuroscience.2006.03.077

